# Optogenetic closed-loop feedback control of the unfolded protein response optimizes protein production

**DOI:** 10.1101/2022.10.27.514034

**Authors:** Moritz Benisch, Dirk Benzinger, Sant Kumar, Hanrong Hu, Mustafa Khammash

## Abstract

In biotechnological protein production processes, the onset of protein unfolding at high gene expression levels leads to diminishing production yields and reduced efficiency. Here we show that *in silico* closed-loop optogenetic feedback control of the unfolded protein response (UPR) clamps gene expression rates at intermediate near-optimal values, leading to significantly improved product titers. Specifically, in a fully-automated custom-built 1L-photobioreactor, we used a cybergenetic control system to steer the level of UPR in yeast to a desired set-point by optogenetically modulating the expression of a hard-to-fold protein based on real-time feedback measurements of the UPR, resulting in 60% higher product titers. This proof-of-concept study paves the way for advanced optimal biotechnology production strategies that diverge from and complement current strategies employing constitutive overexpression or genetically hardwired circuits.

---

*S. cerevisiae* is a widely used organism for recombinant protein production, known for its fast growth kinetics, and its ability to fold and secrete proteins [1]. Many engineering strategies have been applied to further improve the production capacity of yeast and increase protein yield [2]. Intuitively, increasing the rate of gene expression should result in more secreted protein. While this is true at low protein expression rates, maximal expression does not necessarily lead to maximal protein production, but can even reduce protein yield [3] [4]. Consequently, there exists an optimal gene expression rate that maximizes production. The diminishing rate of protein production can be attributed to an overwhelmed cell machinery, resulting in oxidative stress, product misfolding, inclusion bodies, upregulated endoplasmic reticulum associated degradation, and stress-induced genomic instability [5] [6] [7] [8].

The mapping between optimal gene expression rate and optimal protein production is influenced by many different factors, such as growth rate, process stage and complexity of the protein of interest [9]. Thus, a constant, fine-tuned gene expression level [10] is generally only optimal during a limited process window and for a single product. One possible solution is to adjust protein expression automatically based on stress levels using burden-driven, genetic feedback circuits Ceroni et al. [11]. However, for this approach to function optimally, the synthetic circuits themselves need to be fine-tuned to a given product and production condition. Additionally, hard-wired feedback cannot adapt when process requirements change over time.

These challenges can be solved by applying direct *in silico* control on cell internal states similar to what has been established in the industry for process parameters like pH and dissolved oxygen. This allows for the adjustment of feedback strength to the design criteria, ultimately improving protein production. Here we focus our attention on the unfolded protein response (UPR), as it is central to the proper folding of proteins and is often engineered to enhance protein production hinting towards an ideal UPR induction level. Also, it incorporates the aforementioned factors influencing the gene expression to protein yield map (illustrated in figure 1a). We engineered *S. cerevisiae* to optogenetically express a hard-to-fold model protein, as well as a fluorescent sensor of UPR activity. Further, we built a cybergenetic bioreactor platform that automates sampling, dilution, measurement and dynamic real-time feedback of our culture. Ultimately, we study how clamping the mean population stress level to different constant levels affects the final yield of the model protein *α*-amylase.

**Figure 1:**
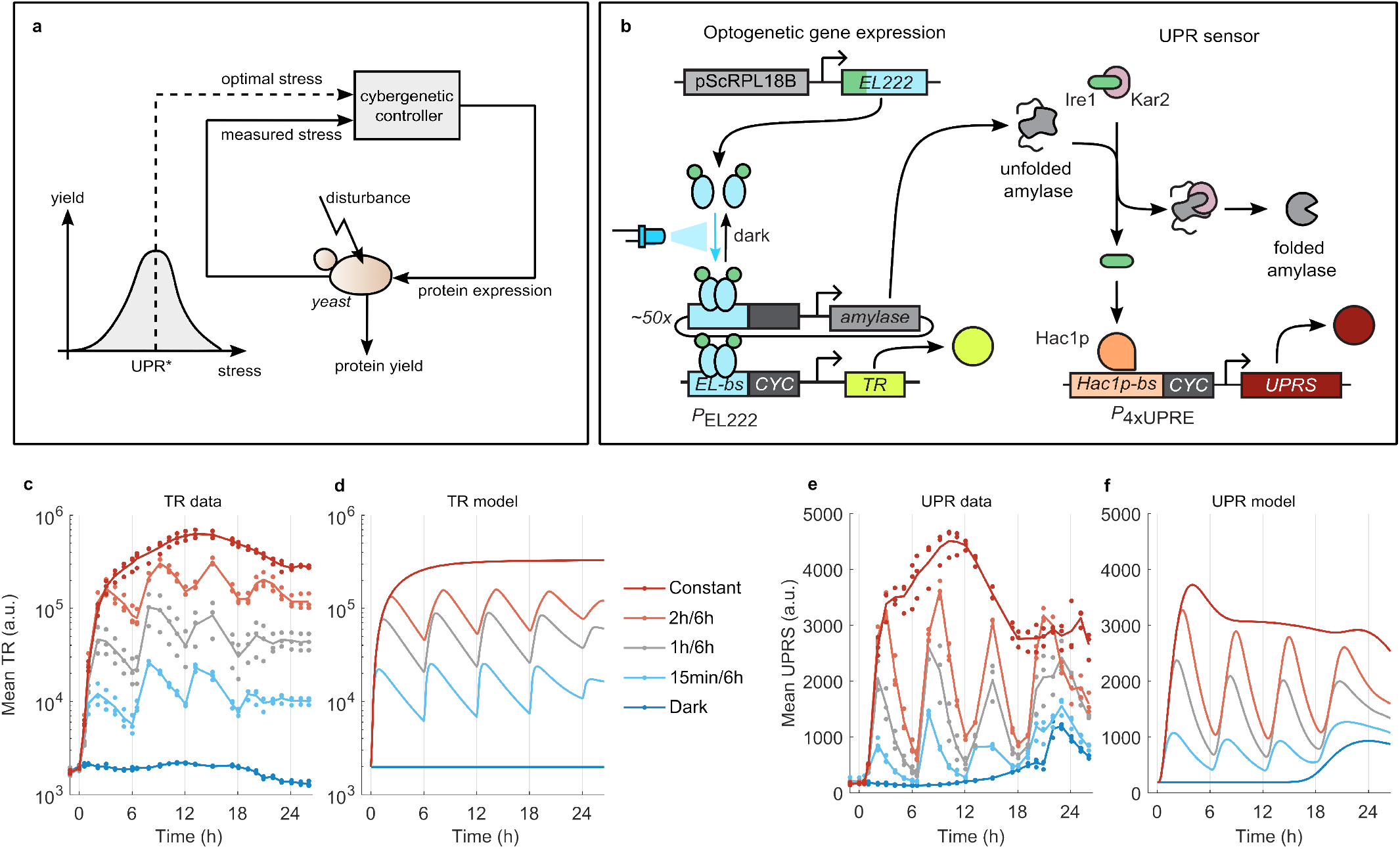
Construction and characterization of a yeast strain for optogenetic protein production and concurrent UPR reporting **a** We postulate that an optimal UPR stress (UPR*) exists which maximizes protein yield. A cybergenetic controller takes this optimal stress input and the measured UPR stress and computes the optimal next protein expression strength. The engineered *S. cerevisiae* produce protein given the protein expression strength and its growth parameters (disturbance). **b** Illustration of the optogenetic expression system and UPR sensor. Constitutive expression of Msn2AD-EL222 from the genome results in Msn2AD-EL222 monomers that reversibly dimerize under blue light and allows binding to the Msn2AD-EL222 binding site (EL-bs) activating transcription from the P_*EL*222_ promoter. P_*EL*222_ drives the expression of *α*-amylase and transcriptional reporter (TR), from a high-copy plasmid and chromosomal integration respectively. Unfolded proteins trigger the unfolded protein response (UPR), which leads to expression of the Hac1p transcription factor, subsequent binding to the P_4*xUPRE*_ and expression of a UPR sensor (UPRS) from a chromosomal integration. **(c-f)** Dynamic response of the mean fluorescence level of TR (**b**) and UPRS (**e**) to different amounts of blue light measured in the water bath setup (n=3, lines represent means of replicates) and compared to the mathematical model fit (**c** and **f**). Cells were grown in the dark, with 15min, 1h, and 2h resp. of light every 6h and with constant light.

This new way of producing proteins using cybergenetics has the potential to revolutionize and massively simplify the upscaling of new proteins of interest. Simultaneous characterization of the production process in combination with closed-loop real-time feedback allows one to reduce overall turnaround times.

In order to implement cybergenetic feedback, it is necessary to be able to control the expression rate of the protein of interest reversibly and in a graded fashion. We engineered *S. cerevisiae* to express the hard-to-fold, secreted protein *α*-amylase under control of an optogenetic transcription factor (TF), consisting of the blue-light-sensitive DNA-binding domain EL222 fused to the Msn2 activation domain (Msn2AD-EL222) [12] [13]. Under blue light, the TF dimerizes, enabling it to bind to its target promoter P_*EL*222_ and thus activate gene expression. In the dark, Msn2AD-EL222 reverts to its transcriptionally inactive state within minutes [12] [13], enabling highly dynamic expression regulation. To dynamically measure the *α*-amylase transcription, we further constructed a transcriptional reporter (referred to as TR) by placing the fluorescent protein Venus under control of P_*EL*222_ (figure 1b). We expect that transcription of *α*-amylase induces the cellular UPR, resulting in the expression of the transcription factor Hac1p. We implemented a UPR sensor (UPRS) by incorporating Hac1p responsive elements [14] in a CYC1 minimal promoter [15] (P_4*xUPRE*_) that drives the expression of the fluorescent protein mScarlet-I (figure 1b).

The media was supplemented with inositol to avoid the triggering of the UPRS by inositol starvation (figure S1). In this supplemented media, we show that global chemical stress results in upregulation of the UPRS (figure S2) and further tested whether the optogenetic expression of *α*-amylase would also trigger the UPR. Figure 1c and 1e show that upon blue light illumination, the *α*-amylase proxy TR is expressed and the UPRS levels follow this rise. This indicates that expression of *α*-amylase induces the UPR. Turning off the blue light stopped the transcription of amylase, reducing the influx of new unfolded proteins to the ER and decreasing the level of UPRS. Light-mediated activation was graded and could be adjusted by increasing the duration of blue light illumination. We developed a mathematical model (fig S5a), which combines the optogenetic activation of gene expression [16], the UPR [17] and growth dynamics (see Supplementary Information). After model fitting, the model captures the dynamics of UPRS and TR well (figure 1d and 1f).

Having established that we can optogenetically control amylase production and follow the resulting UPR activity over time, we sought to investigate how closed-loop control of the UPR affects the overall production of amylase. In order to run closed-loop feedback experiments on a production relevant scale, we developed a custom-built 1L-bioreactor platform which allows for automatic sampling, dilutions, online measurements of cell reporters, proportional-integral control and optogenetic feedback taking inspiration from Milias-Argeitis et al. [18] (figure 2a and S6).

**Figure 2:**
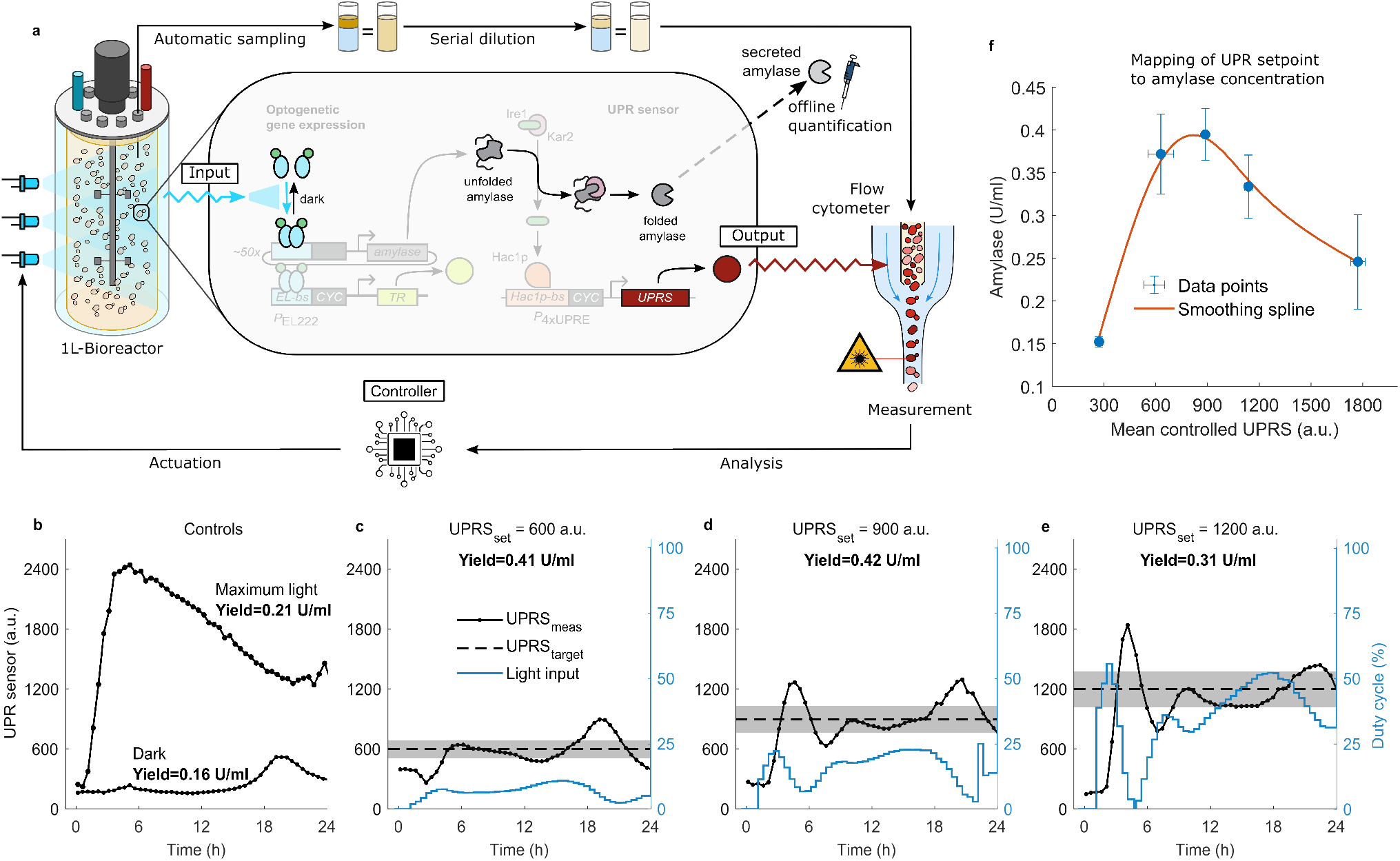
**a** Closed-loop optogenetic 1L-bioreactor platform and engineered *S. cerevisiae*. The bioreactor is placed in a light house to allow illumination of the culture broth. The culture broth is automatically sampled by peristaltic pumps at regular intervals, diluted in two stages with diluent and finally transferred to a flow cytometer for measurement of the cell density and UPRS. The measurements of the UPRS are automatically analyzed, and the PI controller calculates the next light input, closing the feedback loop. **b** Evolution of the UPRS over time for dark and maximum light condition with respective yield of *α*-amylase. **c-e** Evolution of the UPRS over time in the bioreactor setup in the first 24h during which feedback control is performed. Black solid line represents the measured UPRS, dashed the target UPRS level with 15% area around it. The blue line represents the blue light duty cycle in % resulting from closed-loop control. The closed-loop UPRS setpoints are 600 (**c**), 900 (**d**) and 1200 a.u. (**e**) resp. **f** Mean and standard deviation of the final amylase concentration after 70 hours is plotted against the mean and standard deviation of the UPRS during the controlled time (first 24 hours) as shown for panels b-e (two independent experiments per condition, three for maximum light). We perform Welch’s t-test and find that the *α*-amylase yield at the closed-loop setpoints 600 and 900 a.u. is higher than the yield with maximal induction at a significance level of 4.6% and 1.5%. The closed-loop setpoint at 1200 a.u. achieves a yield that can not be deemed statistically higher from maximal induction (*α* = 6.2%).

Using this setup, we first evaluated UPR activity during a batch process without and with constant, maximum blue-light induction, respectively (figure 2b). Without light, cells express a basal level of UPRS throughout the run, with a transient, stationary-phase induced increase after 18 hours. Blue-light induction results in a fast and sharp increase of UPR activity which reaches its maximum at 6 h post-induction, after which the UPR adapts to an intermediate activity. The dynamic behaviour of the UPR highlights the need for closed-loop control to achieve constant levels of activity during a run. The experiment is stopped after 70 hours as the dissolved oxygen plateaus (figure S11).

Next, to evaluate the relation between UPR activity and protein production, we aimed to clamp the UPR to three different levels by optogenetic feedback control. Due to the non-linearity of light intensity to gene expression, we chose to modulate the light input by altering the duty cycle at constant light intensity [16]. Due to the onset of the stationary phase we decided to limit the control to the first 24h and set the light duty cycle to 100% for the next 46 hours (see figure S7 for full time line) [19]. Thus tracking of the desired UPRS value needs to be achieved in a relatively short amount of time, resulting in the need to tune control parameters of the PI-feedback accordingly. To avoid time-intensive experimental parameter tuning, we leveraged our mathematical model to predict the performance of proportional-integral (PI) control parameters (fig S5b) for the closed-loop experiments. Performing closed-loop feedback experiments with these parameters, we achieved tracking of all three setpoints (600, 900, and 1200 a.u. UPRS). This tracking is maintained, after an over- and undershoot, while cells are growing exponentially (100-fold range of cell density, fig. S10). To our knowledge, this is the first implementation of closed-loop feedback of a cell state in a high cell density culture.

We collected the supernatant after 70 hours and measured the *α*-amylase concentration. Figure 2f shows the final amylase concentration vs. the mean controlled UPRS in the first 24 hours after inoculation. One can observe that at higher levels of UPRS (constant illumination), the final *α*-amylase concentration is lower than for the intermediate UPRS setpoints that were achieved using the closed-loop control setup. This follows our hypothesis from figure 1a. For optimal protein production, the cells should be kept at 900 a.u. of UPRS to improve the titer of *α*-amylase by 60% over maximally inducing the cells as well as an increase of 159% over cells grown in the dark. We hypothesize that the cells are running into a limitation of the protein production machinery, most likely in the folding machinery. There is additional evidence, that strong and consistent induction of the UPR over a long time, will lead to the formation of inclusion bodies and trigger the ER-associated degradation machinery [20].

We presented the development of a biological strain that allows optogenetic expression of a model protein in combination with fluorescent reporters about the transcriptional and UPR state of the cell. The presented biological circuit with reversible optogenetic induction of the UPR could be a valuable tool to study the time-course dynamics of the UPR *in vivo*. We initially characterized this circuit in a low volume water bath setup and used the data to fit a mathematical model. This mathematical model predicted control parameters that were used without further fitting to upscale the volume 100-fold. Furthermore, we developed a platform for optogenetic closed-loop feedback control in high cell density cultivations (HCDC). Lastly, we demonstrated that closed-loop feedback control of UPRS allows tight tracking of setpoints during exponential growth, and that an optimal UPR setpoint exists, for which the amount of *α*-amylase is maximal. This implementation of *in silico* closed-loop feedback at a HCDC bioreactor level opens new possibilities about how to express and produce proteins efficiently. Future research could explore more complex induction profiles.

## Acknowledgements

We thank Dr. Stephanie Aoki for help with strain design and useful discussions, Dr. Gregor Schmidt for helpful discussion, optimization of the *α*-amylase quantification and automation of the measurement and Peter Buchmann for the building of the light housing for the optogenetic bioreactor platform. We also thank Dr. Stephanie Aoki and Dr. Joaqeuín Gutiérrez Mena for proofreading the manuscript. This project has received funding from the European Research Council (ERC) under the European Union’s Horizon 2020 research and innovation programme (CyberGenetics; grant agreement 743269).

## Author contributions

M.B., D.B. and M.K. conceptualized the study and wrote the manuscript. M.B. built the engineered *S. cerevisiae* strains, performed the characterization, closed-loop experiments and performed the data analysis. M.B. and H.H. developed the initial modeling framework and performed the computational modeling and optimization of PI gains. M.B. and S.K. developed the automated sampling setup. D.B. performed preliminary experiments on optogenetic amylase production and co-supervised the project. M.K. secured funding and supervised the project.

## S1 Methods

### S1.1 Media

YPD and SD-URA medium for transformations was prepared according to the Clontech Yeast Protocols Handbook [1]. The YPE medium contains 10 g/L yeast extract, 20 g/L peptone and 25 ml/L ethanol. Just before usage, acetaldehyde is added to YPE at a concentration of 0.01 % to reduce the lag time in growth observed otherwise.

Wittrup and Benig [2] developed a media for heterologous protein secretion in *S. cerevisiae*. This SD-2xSCAA medium is used for all precultures and experiments and contains 20 g/L D-glucose, 6.9 g/L yeast nitrogen base without amino acids, folic acid and riboflavin, 1 g/L bovine serum albumin (BSA), amino acids as listed in supplementary table S1, 5.4 g/L Na_2_PO_4_ and 8.56 g/L NaH_2_PO_4_ · H_2_O (pH = 6.0 by H_2_SO_4_). For experiments in the bioreactor, the sodium phosphate buffer salts are replaced with 2 g/l KH_2_PO_4_ and 500 *µl/l* of 20 % PPG P 2’000 is added as antifoam. To prepare SD or YPD agar plates, 20 g/L agar is added to the medium. For SD-2xSCAA plates, BSA was not added.

While running long-term experiments of cells grown in the dark, we observed a sudden increase in UPRS levels after 18 hours for an inositol concentration of 2 mg/l (figure S1). This is the original concentration of inositol in the SD-2xSCAA protein production media [2] [3]. Leber et al. [4] reported that inositol starvation triggers the UPR in yeast while Henry et al. [5] observed that normal SD media is limited in inositol. We thus tested whether the SD media supplemented with synthetic casamino acids was also limited in inositol and supplemented the media with additional myo-inositol to initial concentrations of 2-200 mg/l. Figure S1 shows that supplementation of the media with inositol delayed the onset of UPR induction as well as decreased the induction fold change. Given these results we supplemented our media to a final inositol concentration of 20mg/l.

For precultures and characterization experiments, the antibiotics Hygromycin B and Geneticin G418 were added at a final concentration of 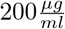. For transformations with Zeocin resistance YPD plates were poured with Zeocin at a concentration of 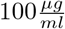. Zeocin was not added during experiments as it is highly unstable when exposed to light.

**Figure S1:**
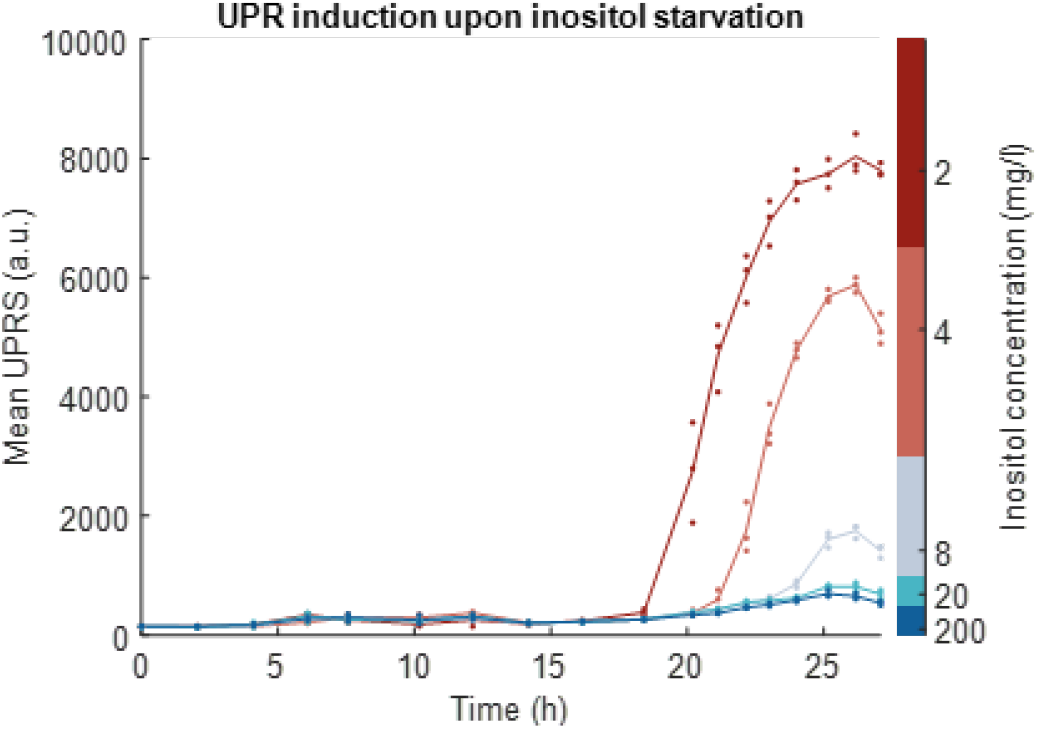
UPR induction upon inositol starvation is observed at different times for different initial concentrations of inositol in culture tubes (n=3). The initial SD-2xSCAA media contains 2mg/l of inositol and was supplemented with myo-inositol to reach the inositol concentrations given on the right.

Amino acids, *α*-amylase, D-glucose, phosphate buffers, myo-inositol and BSA were purchased from Sigma-Aldrich. Geneticin was purchased from Gibco, Zeocin and Hygromycin B from Invitrogen, agar, yeast extract, peptone from BD, yeast nitrogen base from Formedium, ammonium sulfate from Roth and PPG from Fluka.

### S1.2 Culture conditions

All precultures and experiments were grown at 30°C in SD-2xSCAA. Experiments were performed at three different volume scales. Experiments for initial characterization of the UPRS and experiments to determine the ideal concentration of inositol were performed in 10ml culture tubes in 3ml media in a shaking incubator (New Brunswick). Optogenetic characterization experiments were performed in a water bath setup [6]. Vials can be individually illuminated with a blue LED (*λ* = 450*nm*) below each vial. A stir bar is added to each tube and the 4ml culture is stirred at 900 rpm. For the final amylase production, cells were grown in a 1.0l Eppendorf BioFlo120 water-jacketed bioreactor with a working volume of 500 ml, 300 rpm agitation and 2.0 SLPM (standard liter per minute). The dissolved oxygen was controlled above 20% using the InPro 6800 Series O_2_ Sensor (Mettler Toledo, Switzerland) by automatic adjustment of the impeller speed (between 300 and 500 rpm) and air flow rate (between 2.0 and 3.0 SLPM). The pH was maintained at between 5.95 and 6.05 by a Type 405-DPAS-SC pH sensor (Mettler Toledo, Switzerland) using 4 M NaOH and 10% H_2_SO_4_. All fermentations were done in biological duplicates (figure S8 and S9).

The bioreactor unit is placed in a custom-built light housing that protects it from any external light source present in the laboratory. Six Adafruit NeoPixel NeoMatrix LED panels with 8×8 LEDs were installed at the side of a light housing to illuminate the bioreactor. As continuous illumination of the bioreactor with the panels resulted in heating up of the pads, heat sinks were installed on the back of the LED panels to passively cool them. All process parameters can be extracted, visualized and changed through a custom-build MATLAB script.

MATLAB triggers the sampling every 30 minutes, while two Arduino Unos are executing the pumping steps. Samples are pumped automatically every 30 minutes through the sampling port of the bioreactor. The sample is then separated from the rest of the bioreactor media by air and pushed towards a first dilution stage. The sample remaining in the tube from bioreactor to the first dilution stage is pushed back into the bioreactor container maintaining sterile conditions for the culture broth. The collected sample is diluted with the help of a second pump and mixed in the process. This process is repeated in a second dilution stage to achieve dilutions ranging between 1- and 400-fold to cell densities that allow for measurements in a flow cytometer 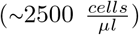. The diluted sample is pumped to a sampling vial of a flow cytometer and measurements are automatically acquired through an application programming interface (API), exported and analyzed. The PI controller calculates the next light input and an Arduino Mega changes the light input accordingly. After acquisition, the diluted samples are pumped into a waste container and the vials are washed again with more diluent. More details can be found in figure S6. This experimental platform is to our knowledge the first platform for closed-loop optogenetic feedback control at this size and cell density [7].

For all experiments, yeast strains were streaked from the frozen glycerol stock on a SD-2xSCAA plate and incubated at 30°C for 3 days. Precultures were inoculated from single colonies into 10 ml (for characterization) or 25 mL (for bioreactor experiments) of SD-2xSCAA media in a 125 or respectively 250 mL baffled flask and incubated overnight to an *OD*_600_ ≈ 1.25. The preculture cell density was measured by flow cytometry and experiments started at a cell density of 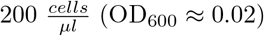. All experiments were shielded from ambient light to avoid unwanted optogenetic activation of the circuit.

**Table S1:** Amino acid concentration for SD-2xSCAA.

| Amino acid | Concentration (mg/l) |
| --- | --- |
| Arginine | 190 |
| Aspartic Acid | 400 |
| Glutamic acid | 1260 |
| Glycine | 130 |
| Histidine | 140 |
| Isoleucin | 290 |
| Leucine | 400 |
| Lysine | 440 |
| Methionine | 108 |
| Phenylalanine | 200 |
| Threonine | 220 |
| Tryptophan | 40 |
| Tyrosine | 52 |
| Valine | 380 |

### S1.3 Flow cytometry

The fluorescence and cell density measurements were performed on a Beckman Coulter CytoFLEX S. The internal quality control program was run before experiments using CytoFLEX Daily QC Fluorospheres. Events were gated in the FSC-A and SSC-A channels to only measure single cell events. Samples were diluted with PBS to cell densities of 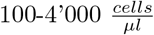. All fluorescence measurements are reported as the mean of the gated population in arbitrary units (a.u.).

### S1.4 Amylase quantification procedure

We used the Megazyme alpha-amylase assay kit to measure the concentration of alpha-amylase in the supernatant. Commercial *α*-amylase (Sigma-Aldrich) from *Aspergillus oryzae* (Cat. Nr. 10065) was used as a standard. 2 ml of sample was centrifuged at 4’000 rcf for 5 minutes to separate cells from the amylase containing supernatant. 1 ml of supernatant was mixed with 10 *µl* of 11mM sodium azide to stop microbial growth. The Megazyme alpha-amylase assay reaction volume was scaled-down from 3.4 ml total volume to 250 *µl* total volume to enable automation and quantification in 96-well plates. The assay was further performed at room temperature, to reduce the amount of evaporation in the 96-well plate and the incubation time with HR reagent extended to 15 minutes. The following steps for quantification were performed. 50 *µl* of biologically inactivated supernatant was transferred into a 96-well plate and incubated at room temperature for 5 minutes. 50 *µl* of HR solution was added to each sample to start the reaction. The plate was shaken at 1’000 rpm for 15 minutes before adding 150 *µl* of 40 g/l Na_3_PO_4_ (pH=11). The stop buffer was prepared by dissolving 92.768 g of Na_3_PO_4_ · 12 H_2_O in 1l water. The pipetting was performed on a Hamilton microlab star and measurement of the absorbance at 400nm with a Tecan Infinite M200Pro.

### S1.5 Strains and plasmids

The genotype of the background, all further strains and plasmids is shown in supplementary table S2 and S3. The phototoxicity experiments (figure S3) were performed with yMB9, all other experiments with yMB44. The yeast production strain *S. cerevisiae* Cen.PK 530-1C (kindly provided by Jens Nielsen, Chalmers University of Technology, Sweden) was used for all work in this publication. Cen.PK 530-1C has a triose phosphate isomerase (tpi1) deletion and is thus unable to metabolize glucose and can only grow on ethanol. The high-copy production plasmid pAlphaAmyCPOT (kindly provided by Jens Nielsen, Chalmers University of Technology, Sweden [8]) contains a *S. pombe* POT1 expression cassette, enabling metabolization of glucose in the tpi1-deletion background, thus allowing for selection of plasmid retention by using glucose as carbon source. The *α*-amylase expression cassette consisting of the P_*EL*222_ promoter, the amylase gene and the TPI1 terminator was placed on this plasmid. The Msn2AD-EL222 expression cassette was expressed from the constitutive ScRPL18B promoter and integrated in the URA3 locus by uracil selection. The transcriptional reporter consists of the yellow fluorescent protein Venus [9] tagged with an N-degron Ubiquitin (Ubi-Y, [10]), under the control of the P_*EL*222_ promoter and was genomically integrated in the HO locus and selected for by Hygromycin B. We implemented a UPR sensor (UPRS) by placing mScarlet-I [11] tagged with an N-degron Ubiquitin (Ubi-Y, [10]), under the control P_4*xUPRE*_).

The UPRS was integrated in the YIRC∆6 locus [12] and selected for by Zeocin. We tested the functionality of the UPRS by using dithiothreitol (DTT), which is a global reducing agent that breaks intramolecular disulfide bonds, leading to protein unfolding and activation of the UPR. Figure S2 shows a graded response of the UPRS to DTT concentrations between 0.2 and 2 mM and changes in the reporter levels can be observed 1 hour after the addition of DTT.

**Figure S2:**
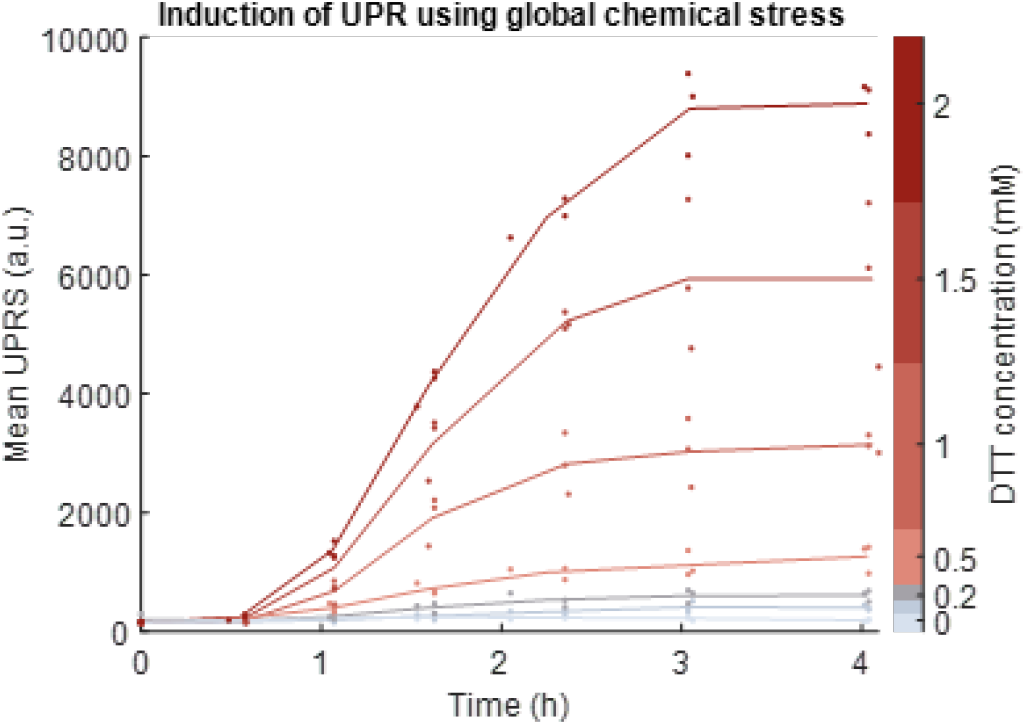
Graded induction of the UPR using the global chemical stressor dithiothreitol (DTT) at different concentrations over time in culture tubes (n=3). DTT is a global reducing agent that breaks intramolecular disulfide bonds, leading to protein unfolding and activation of the UPR. Lines represent means of replicates.

We performed further controls to ensure that UPRS is not triggered through phototoxicity. To this end, we integrated the UPRS in a strain lacking the amylase expression plasmid and tested whether the UPRS would sense induction by DTT and light. Figure S3 shows that only induction with DTT triggered a detectable change in UPRS expression. Shining blue light on cells results in no detectable difference in UPRS compared to cells grown in the dark.

Top 10 competent *E. coli* was used for all molecular cloning. Plasmid construction was done using standard molecular cloning techniques or using the yeast molecular cloning toolkit (YTK) from Lee et al. [10]. Yeast transformations were performed according to the protocol from Gietz and Schiestl [13]. For genomic integrations plasmids were cut with Not-I HF (New England Biolabs) and purified using with a DNA Clean & Concentrator Kit (Zymo Research). Successful transformation was checked with the genotyping protocol from Lõoke et al. [14].

**Figure S3:**
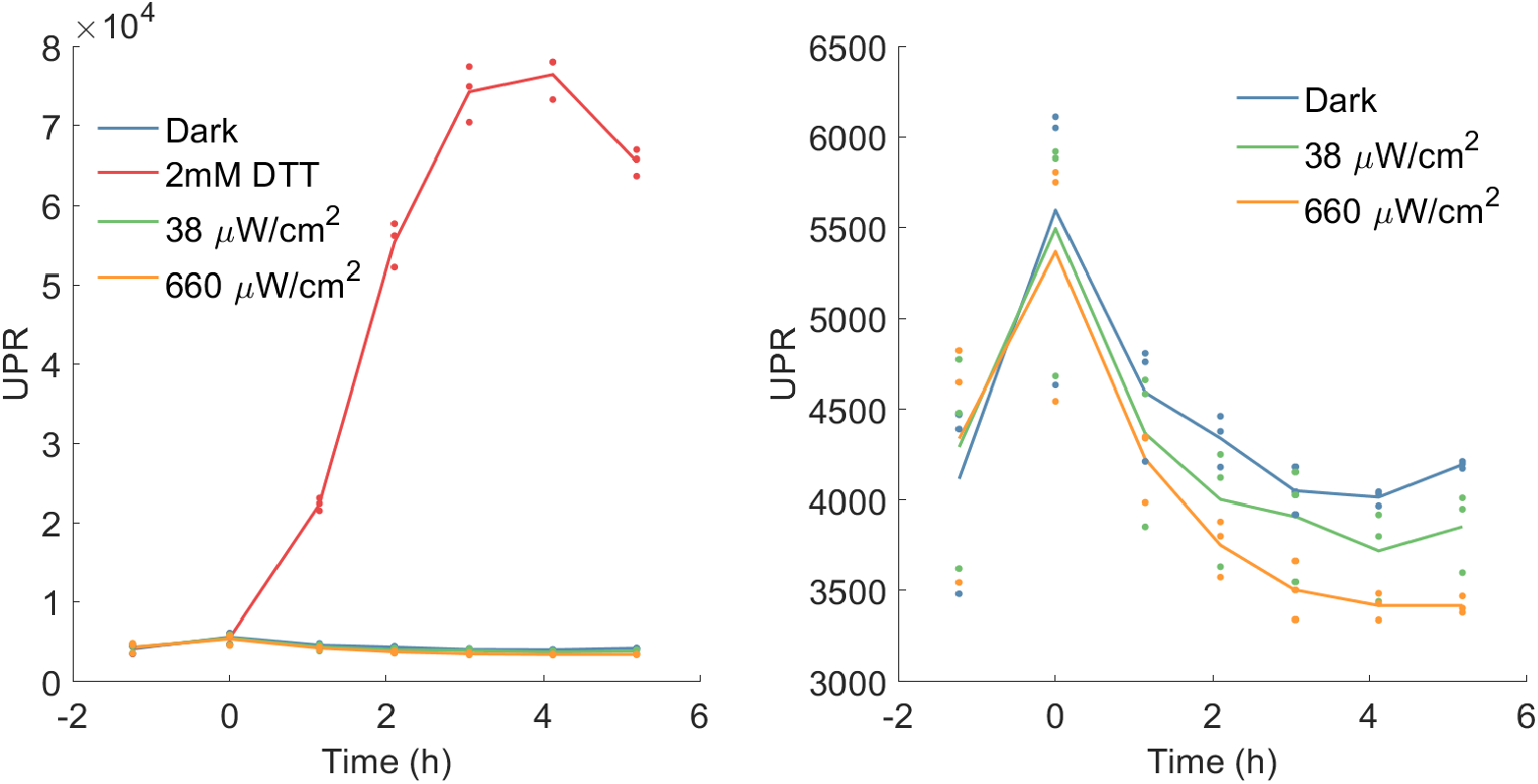
Evolution of the unfolded protein response sensor over time in a strain lacking the *α*-amylase gene expression cassette (yMB9) in response to different inducers. Cells were induced with 2mM DTT or blue light (38 and 660 *µW/cm*^2^) at *t*_0_ = 0*h*. (left) All conditions, with a clear induction of the UPRS caused by DTT and basal expression for cells grown in the dark and exposed to light. (right) Zoom to the three lower conditions with little observable difference of the three conditions. The fluorescent protein used for the UPRS is Venus and so the values are not comparable to the UPRS used in the other experiments. (n=3)

**Table S2:** Plasmids used for strain construction. Promoters are represented by “pr”, terminators are represented by “t”.

| Plasmid | Description | Marker | Source |
| --- | --- | --- | --- |
| pAlphaAmyCPOT | TPIpr-Amylase | POT1 | [8] |
| pYMB4 | 4xUPREpr-UbiY-Venus-ScPGK1t | Hygromycin B | This study |
| pYMB10 | EL222pr-Amylase-TPI1t | POT1 | This study |
| pYMB12v | EL222pr-UbiY-Venus-ScPGK1t | Hygromycin B | This study |
| pYMB15m | 4xUPREpr-UbiY-mScarletI-ScPGK1t | Zeocin | This study |
| pYTKmk48 | ScRPL18Bpr-Msn2AD-EL222-ScENO2t | URA | [15] |

**Table S3:** Strains used in this study.

| Name | Genotype | Source | Data shown in figure |
| --- | --- | --- | --- |
| CEN.PK 530-1D | MATα ura3-52 HIS3 LEU2 TRP1 SUC2 MAL2-8<sup>c</sup> tpi1(41-707)::loxP-KanMX4-loxP | [8] | - |
| CEN.PK 113-7D | MATα URA3 HIS3 LEU2 TRP1 SUC2 MAL2-8<sup>c</sup> | [8] | - |
| yMB9 | CEN.PK 113-7D, HO::pYMB4 | This study | S3 |
| yMB31 | CEN.PK 530-1D, pYMB10 | This study | - |
| yMB32 | yMB32, URA3::pYTKmk48 | This study | - |
| yMB39 | yMB32, HO::pYMB12v | This study | - |
| yMB44 | yMB39, YIRCΔ6IX::pYMB15m | This study | 1, S5, 2, S4, S7, S8, S9, S10 |

### S1.6 Mathematical model

The mathematical model consists of two modules describing Msn2AD-EL222-controlled gene expression regulation and UPR dynamics, respectively (see fig. S5a).

The first module consists of three ordinary differential equations (ODEs) (previously described in [6]) modelling the upstream optogenetic circuit with activation of the transcription factor Msn2AD-EL222 (S1) and Msn2AD-EL222 dependent mRNA expression (S2). Additionally, the translation of the transcriptional reporter from mRNA is modelled (S4). In more detail, blue light input (*I*_*eff*_) triggers structural changes and homodimerization of the transcription factor Msn2AD-EL222 (TF) (S1). The intensity between 0% and 100% is scaled by 0.0019. We assume that the total amount of transcription factor TF_tot_ and the rate of monomerization *k*_*off*_ are both constant. Promoter binding of the transcription factor is fast, so that the rate of mRNA transcription is a direct function of TF, *n* and *K*_*m*_ with *k*_*m*_ as a scaling factor (S2). The degradation rate of mRNA is the basal degradation rate plus dilution from growth (S3). The fluorescent transcriptional reporter (TR, S4) is translated from mRNA with translation rate *k*_*u,trans*_. After initial fitting the values for *k*_*m,basal*_ and *n* were close to 0 resp. 1. To avoid overfitting, these parameters were not fitted but kept at 0 and 1.

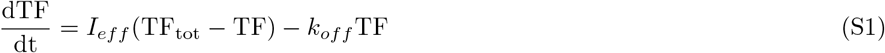

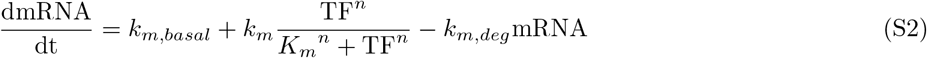

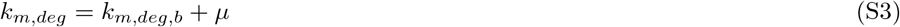

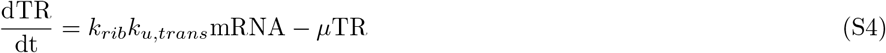

The second module describes the dynamics of the UPR and is based on the minimal model by Trusina et al. [16]. It consists of mRNA translation to unfolded proteins (U, S5a), activation of Hac1p (H) by U (S6) and the resulting upregulation of chaperones, oxidoreductases, glycosylating enzymes and ER degradation components (all combined in node C, S7). C reduces the number of unfolded proteins in the ER, by either folding, secreting the proteins or degrading it and at the same time inhibits H (S5b). The UPRS is transcribed as H binds to the UPR specific promoter (S9).

In the original UPR model [16] the formation of new unfolded proteins U is modelled through a switch-like stress term or a constant translation rate of new polypeptides into the ER. We connect these two modules by using the time-varying translation rate of mRNA (*k*_*u,trans*_mRNA) from our optogenetic module as input to the second module (i.e. the production rate of unfolded proteins (U, S5a).

Upon presence of U, Ire1 clusters on the ER surface. Due to the proximity of Ire1, they transautophosphorylates each other resulting in an activation of the endoribonuclease function of Ire1 (*Ire*1_*act*_, S8). This endoribonuclease excises an intron of the unspliced Hac1 mRNA [17]. The translation process of now spliced Hac1 mRNA is then initiated to produce the potent transcription factor Hac1p (H, S6). The minimal model reduces the splicing and translation steps of Hac1 mRNA into one ODE (equation S6), assuming that at any time point Hac1p protein concentration is proportional to Hac1 mRNA. The assumption is valid because the half-life of Hac1 protein (∼ 1.5min, [18]) is much shorter than Hac1 mRNA (∼ 30min)([17] [19]). The basal degradation rate of H was assumed to follow mRNA kinetics, because of the short half-life. The model describes the regulation of the UPR with strength parameters *α, β, γ* and *δ* (equation S5, S6 and S7).

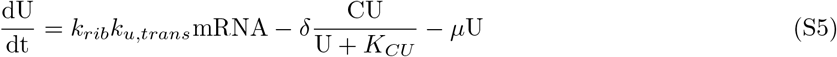

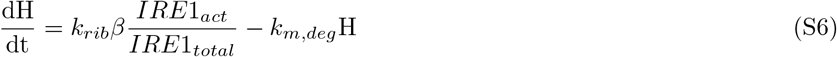

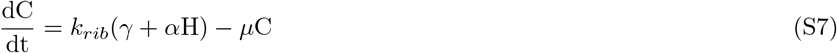

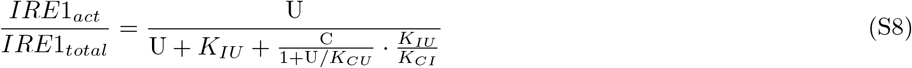

For the UPRS, we assume a linear activation from H to UPRS with strength *k*_*UPR,trans*_ and degradation rate *k*_*UPRS,deg*_ (S9). While supplementation of the media with additional inositol reduced the amount of induction of the UPRS (see figure 1d), still some activation of the UPRS at the end of exponential growth was observed. To capture this induction, a substrate dependent production term of UPRS was introduced (*k*_*UPRs,stat*_, S10).

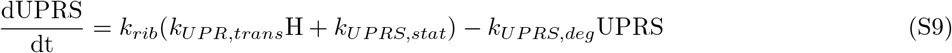

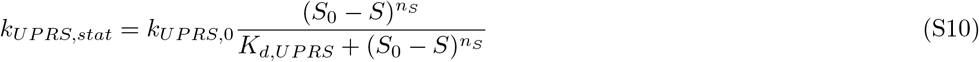

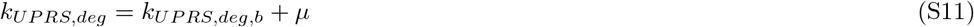

The evolution of the cell density X and limiting substrate concentration S is modelled with equations S12 and S13. Production of biomass X leads to the depletion of the substrate S with a yield factor *Y*_*X/S*_.

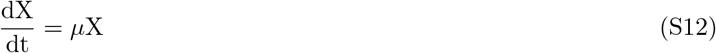

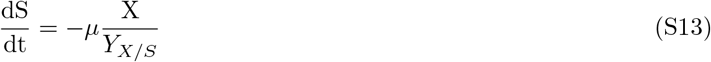

The growth rate *µ* is described through Monods law [20] with a limiting substrate S. We observed the impact of a high transcriptional load on growth rate (figure S4). Cells which are illuminated with maximum light and light pulses of 2h duration every 6h grew slower than cells in the dark. We thus chose to use an inhibitory factor in the growth law of *µ* that incorporates the dose-dependent growth reduction caused by the presence of unfolded proteins (S14). The hypothesis is that the cell growth rate is primarily affected by unfold protein U. Equation S14 describes the burden to cell growth, where *µ*_*max*_ and *U*_*max*_ correspond to the maximal cell growth rate and maximal amount of unfolded protein that the optogenetic circuit can induce. Apart from the burden induced by UPR, factors such as the toxicity of fluorescent reporters are not considered here.

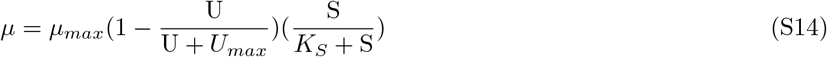

Metzl-Raz et al. [21] show that the ribosomal fraction correlates with the growth rate. While the cells are growing exponentially the fraction of ribosomes is higher, than when cells are going into stationary phase. We see this effect as well, where exponentially growing cells are highly expressing the transcriptional reporter, whereas the amount of TR reduces as cells are in stationarity (see e.g. figure S7 and S9). To capture this behaviour all translation rates are multiplied with the factor *k*_*rib*_ (S15).

**Figure S4:**
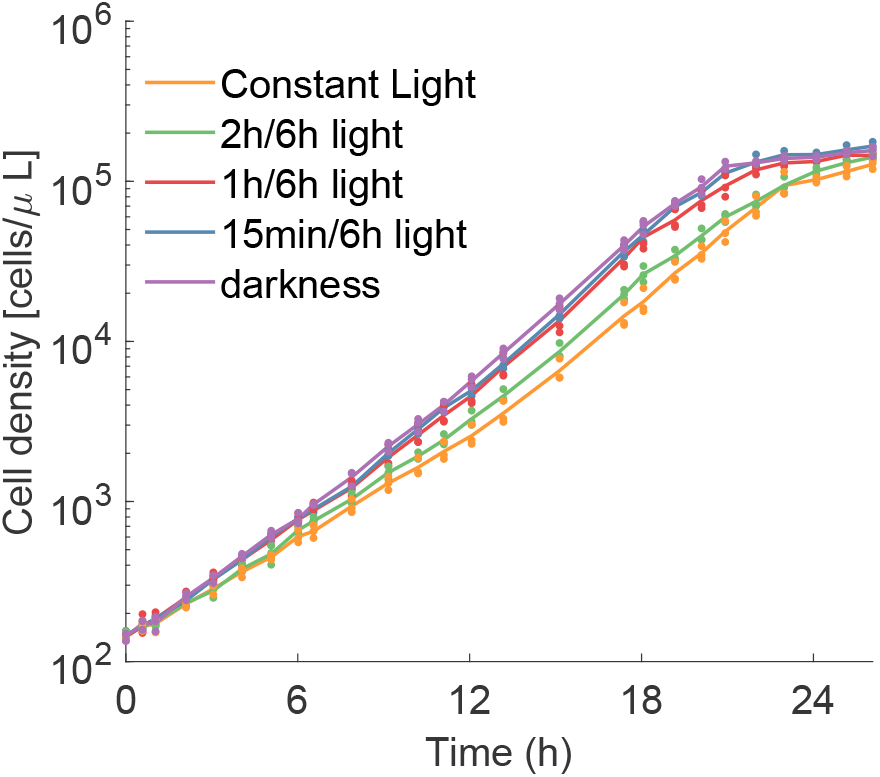
Cell density as measured by flow cytometry in cells/*µ*l over time for the five conditions shown in figure 1. The impact of a high translational load on growth rate can be observed. Cells which are illuminated with constant maximum light and light pulses of 2h duration every 6h grew slower than cells in the dark. (n=3)

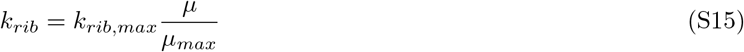

To fit the model to data, experiments for 5 different conditions in triplicates were recorded (fig 1b and 1e). In these experiments, we recorded the fluorescence of TR, UPRS, as well as the cell density in Events/*µl* (fig. S4). The fluorescence data was preprocessed, by subtracting the background fluorescence from the data. The mean of the triplicates was used for fitting. Simulations and model fitting were performed using Matlab (R2020b, Mathworks). The error metric for fitting is calculated by taking the sum of squared errors between model and data of TR, UPR and cell density and normalizing each sum by the mean squared data of TR, UPR and cell density, respectively. This ensures, that the magnitude of the sums is comparable. An additional penalty is applied for the initial point to ensure that the steady state is fitted well. The in-built function fmincon is used for fitting the mathematical model. The final model parameters are listed in supplementary table S4.

### S1.7 Prediction of closed-loop proportional integral control parameters

The mathematical model with fitted parameters is used to predict the performance of proportional integral controllers (PI) with varying control gains K_P_ and K_I_. Figure S5b shows the mean error in the first 18 hours between UPRS setpoint and UPRS as predicted by the model on a grid of 2’500 K_P_ and K_I_ parameter combinations. This PI parameter tuning is repeated for 3 setpoints of the UPRS. Only a limited set of parameter combinations was able to achieve tracking of the setpoint. One can observe that the higher the setpoint, the steeper the resulting surface response. The operating point is chosen to be in close proximity to the lowest region. A design decision was made to pick a slightly higher K_I_, in order to reduce the steady-state error while accepting slightly more oscillatory behaviour.

**Table S4:** Estimated parameters for the optogenetic-UPR model.

| Parameter | Fitted or fixed | Value |
| --- | --- | --- |
| $k_{off}$ | Fitted | 4.77 |
| $TF_{tot}$ | Fitted | $2.23 \cdot 10^3$ |
| $k_{m,basal}$ | Fixed | 0 |
| $k_m$ | Fitted | $2.51 \cdot 10^4$ |
| $n$ | Fixed | 1 |
| $K_m$ | Fitted | $1.50 \cdot 10^3$ |
| $k_{m,deg,b}$ | Fitted | $5.24 \cdot 10^{-2}$ |
| $k_u$ | Fitted | $1.81 \cdot 10^1$ |
| $\delta$ | Fitted | $2.55 \cdot 10^{-2}$ |
| $K_{CU}$ | Fitted | $2.51 \cdot 10^4$ |
| $\beta$ | Fitted | $4.08 \cdot 10^2$ |
| $\gamma$ | Fitted | $9.11 \cdot 10^{-3}$ |
| $\alpha$ | Fitted | $3.75 \cdot 10^1$ |
| $K_{IU}$ | Fitted | $9.93 \cdot 10^5$ |
| $K_{CI}$ | Fitted | $2.55 \cdot 10^3$ |
| $k_{UPR,trans}$ | Fitted | 2.36 |
| $k_{UPRS,0}$ | Fitted | $2.14 \cdot 10^1$ |
| $S_0$ | Fitted | 5.00 |
| $n_S$ | Fitted | 4.20 |
| $K_{d,UPRS}$ | Fitted | 2.60 |
| $k_{UPRS,deg,b}$ | Fitted | $1.73 \cdot 10^{-3}$ |
| $Y_{X/S}$ | Fitted | $3.89 \cdot 10^4$ |
| $\mu_{max}$ | Fitted | $1.47 \cdot 10^{-2}$ |
| $U_{max}$ | Fitted | $4.42 \cdot 10^5$ |
| $K_S$ | Fitted | 8.31 |
| $k_{rib,max}$ | Fitted | 1.04 |

**Figure S5:**
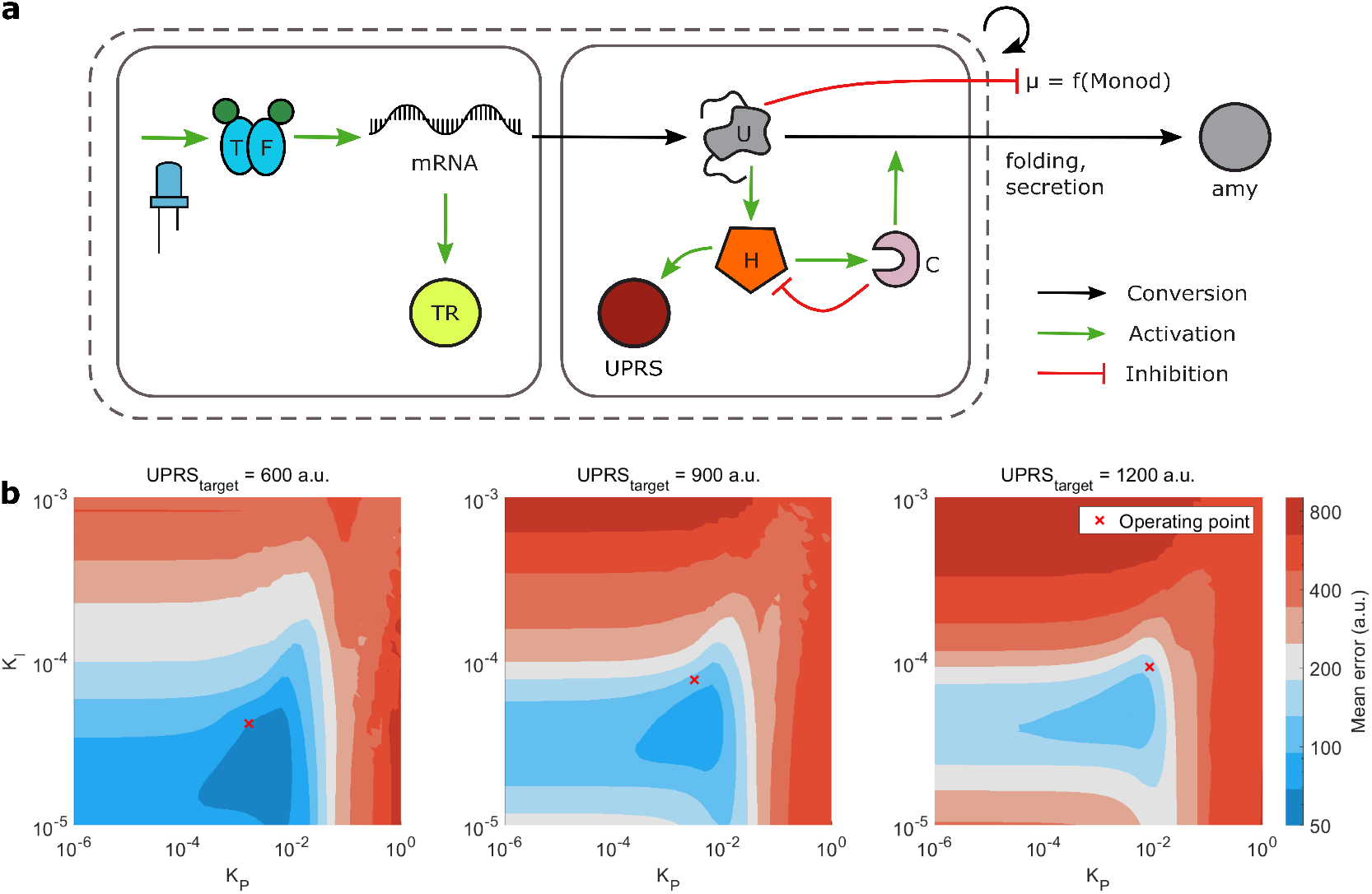
**a**. Structure of the mathematical model, encompassing the description of the optogenetics [6] with activation of the transcription factor TF by light and subsequent transcription to mRNA and translation to TR and unfolded proteins (U). The second module describes the UPR dynamics [16] with activation of Hac1p (H) in the presence of U. H triggers the upregulation of chaperones and related enzymes (C) and UPRS, while C reduces the amount of U by folding, secretion or degradation. **b**. Surface response of the mean error in a.u. to varying values of proportional (K_P_) and integral (K_I_) controller gain for three different target values of the UPRS.

## S2 Supplementary figures

**Figure S6:**
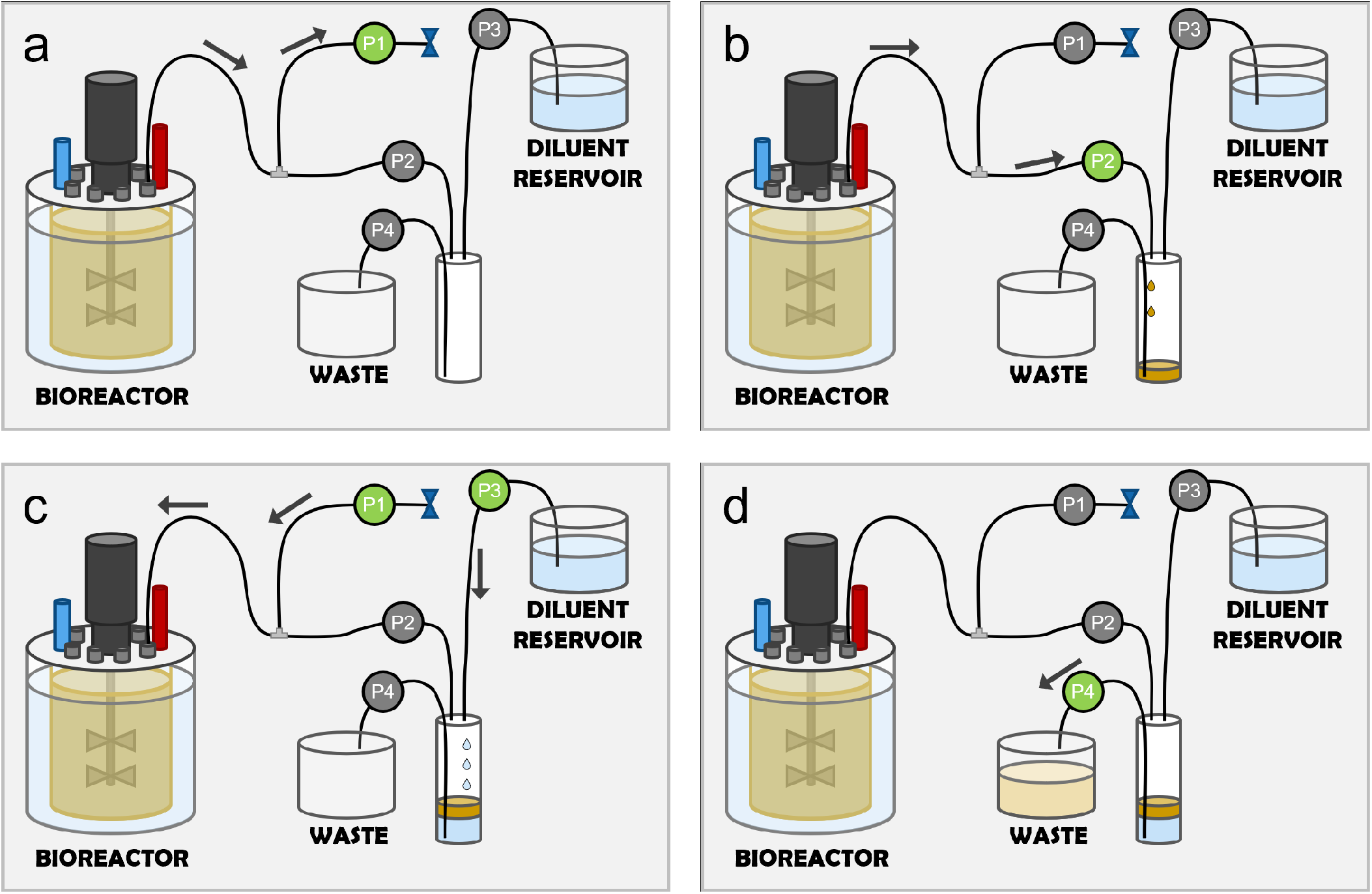
Automatic sampling setup as shown in figure 2 in more detail. In step a, sample is pumped from the bioreactor through a T-piece into tubing of pump 1 (P1). This allows a more precise amount of sample to be transferred through pump 2 (P2) towards the sample vial, as the sample is primed until the T-piece. During this step, the pump direction of P1 is reversed three times to ensure proper mixing of sample in the pump tubing. In step b, sample is pumped through P2 into the sample vial. In step c, the pump direction of P1 is reversed and the sample pushed back into the bioreactor. A sterile filter behind P1 ensures continuous sterility during sampling. At the same time, the sample is diluted with pump 3 (P3). Between steps c and d, sample is transferred on to the next dilution step, similar as in step a from bioreactor to sample vial 1. After transfer, the sample is removed with pump 4 (P4) into a waste reservoir. The vial is rinsed with more diluent (P3) and the diluent removed again with P4. The pumps used for the sampling stem from the Chi.Bio setup and were used for their versatility and low price.

**Figure S7:**
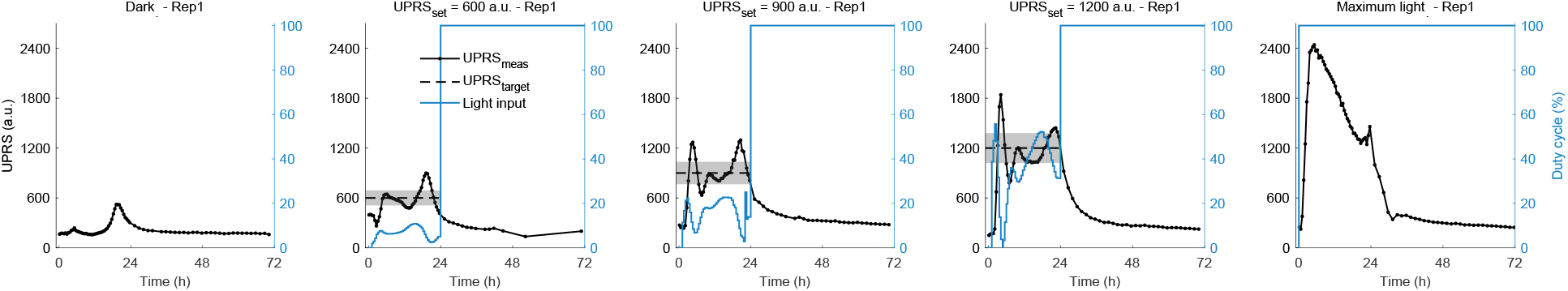
Evolution of UPRS over the entire 72 hours of experiment for repetition 1.

**Figure S8:**
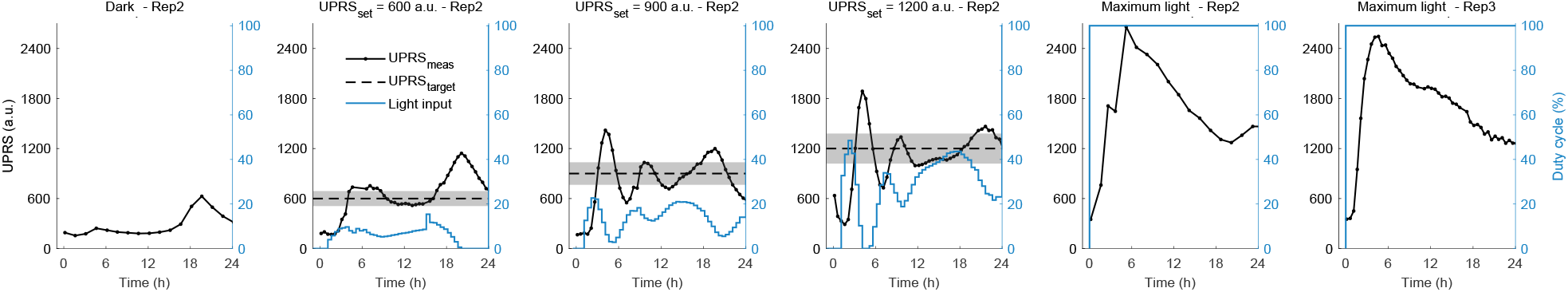
Evolution of UPRS for the 2nd replicate and 3rd replicate for maximum light over the first 24 hours of experiment.

**Figure S9:**
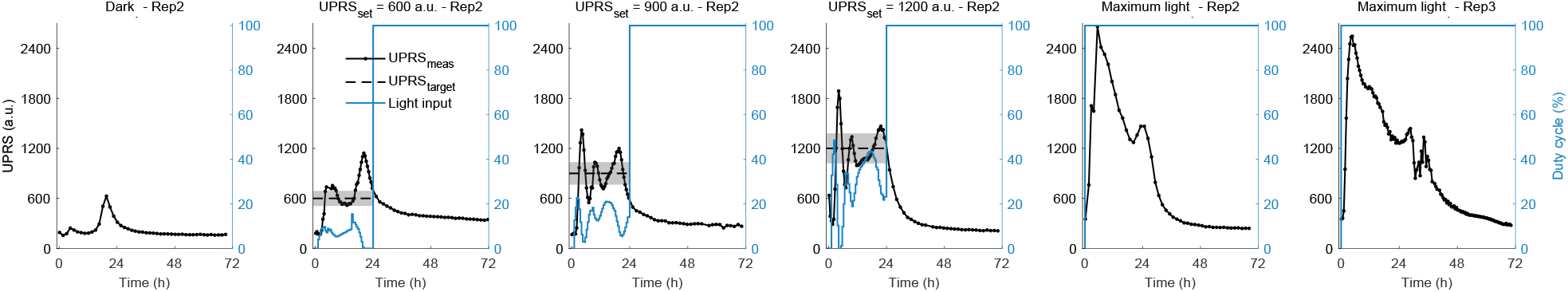
Evolution of UPRS for the 2nd replicate over the full 72 hours of experiment.

**Figure S10:**
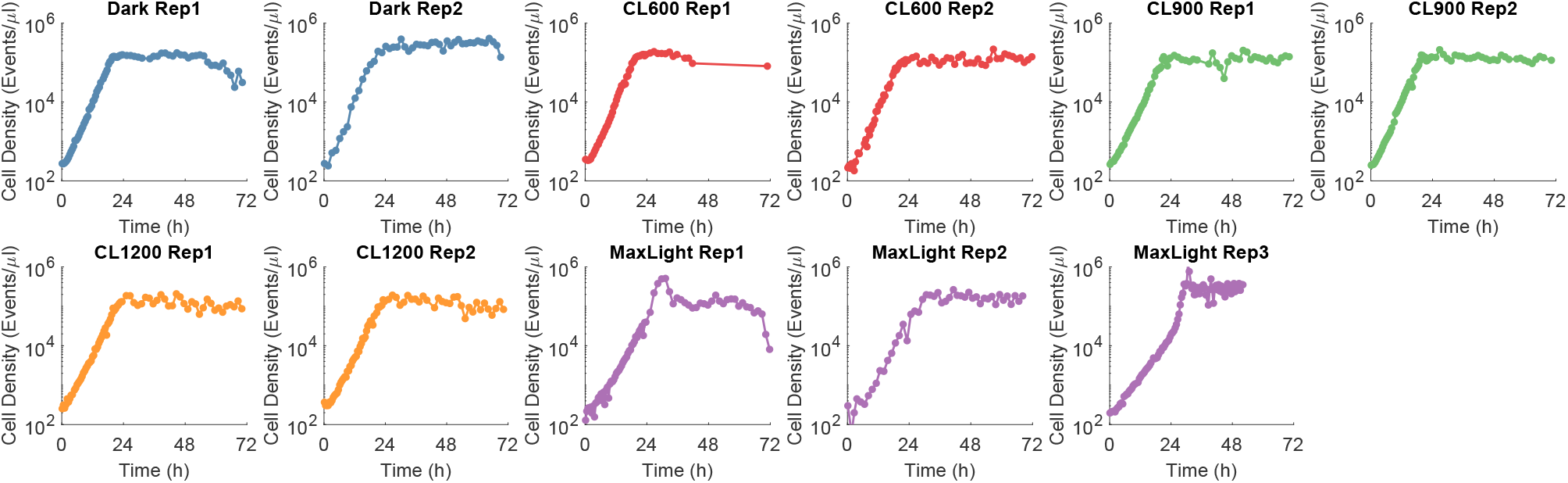
Evolution of the cell density in cells per *µ* l over the full 72 hours of experiment.

**Figure S11:**
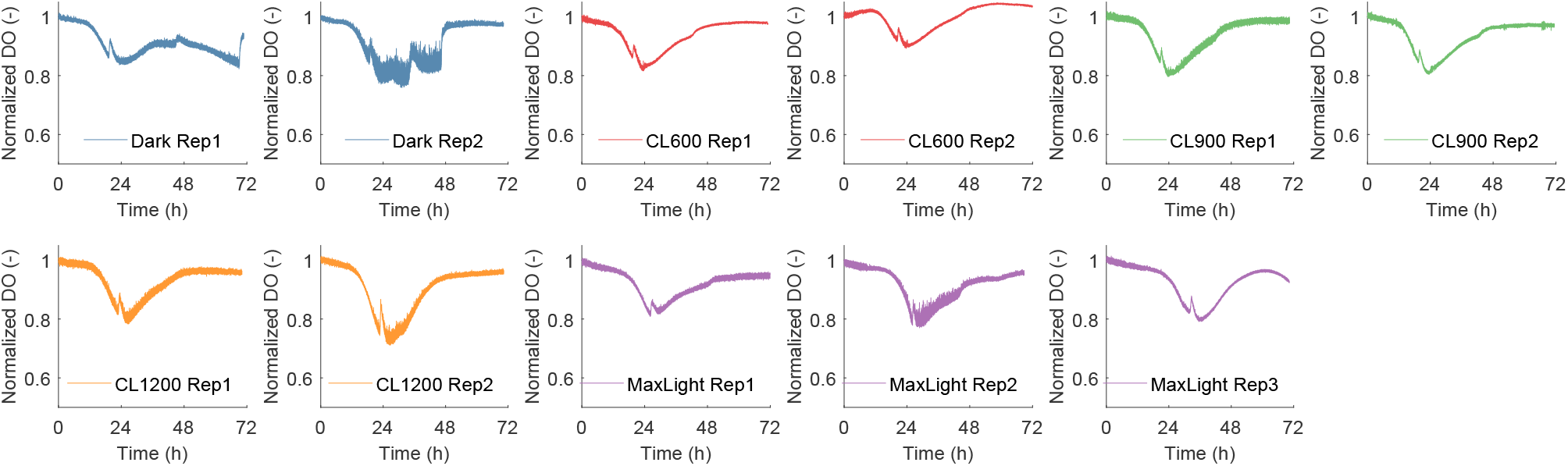
Evolution of the dissolved oxygen (DO) normalized by the initial DO value at t=0h.

